# Fluid Intelligence and Time Management and Artificial Neural Networks

**DOI:** 10.1101/2020.10.29.361410

**Authors:** Markus Ville Tiitto, Robert A. Lodder

## Abstract

Attention deficit hyperactivity disorder (ADHD) is a neurodevelopmental disorder characterized by inattention, hyperactivity, and impulsivity. Our lab is currently conducting a pilot study to assess the effects of the online game Minecraft as a therapeutic video game (TVG) to train executive function deficits in children with ADHD. The effect of the TVG intervention in combination with stimulants is being investigated to develop a drug-device combination therapy that can address both the dopaminergic dysfunction and executive function deficits present in ADHD. Although the results of this study will be used to guide the clinical development process, additional guidance for the optimization of the executive function training activities will be provided by a computational model of executive functions built with artificial neural networks (ANNs). This model uses ANNs to complete virtual tasks resembling the executive function training activities that the study subjects practice in the Minecraft world, and the schedule of virtual tasks that result in maximum improvements in ANN performance on these tasks will be investigated as a method to inform the selection of training regimens in future clinical studies. This study first proposes the use of recurrent neural networks to model the fluid intelligence executive function. This model is then combined with a previously developed model using convolutional neural networks to model working memory and prepotent impulsivity to produce virtual “subjects” who complete a computational simulation of a Time Management task that requires the use of both of these executive functions to complete. The capability of this model to produce groups of virtual “subjects” with significantly different levels of performance on the Time Management task is demonstrated.

## Background

Attention Deficit Hyperactivity Disorder (ADHD) is a highly prevalent neurodevelopmental disorder in children that is characterized by symptoms of inattention, hyperactivity, and impulsivity^1^. ADHD can cause difficulties in academic and social functioning, and can lead to adverse long-term outcomes such as drug addiction and criminal behavior^2^. While the exact cause of ADHD remains unknown, drug therapies have proven successful in controlling symptoms. However, these drug therapies have not proven effective for improving long-term outcomes^2^, their side effects limit use in a significant number of patients, and concerns about their safety has led to a wide preference for non-drug therapies^3^.

An emerging alternative strategy for the treatment of ADHD is the training of executive functions, or a set of conscious mental processes that regulate attention and behavior^4^. Executive function deficits have been proposed as a potential cause for ADHD^5^, and computerized executive function training programs have been investigated as a potential treatment option^6^. The majority of these programs have been designed to target a single core executive function, such as working memory, that is thought to exert far-reaching effects across multiple areas of behavior^4^. While these programs were developed with the expectation that amelioration of these core executive function deficits would lead to improvements in ADHD symptoms and overall functioning, evidence of these far-transfer effects has thus far been limited. Although several studies have shown improvements in parent ratings of ADHD symptoms, teacher ratings and academic performance remained largely unaffected, indicating a lack of generalization of improvements in executive function across multiple environments^6,7^.

Multiple strategies have emerged to address this lack of far-transfer effects resulting from executive function training interventions. Firstly, video game environments have been developed as a setting for executive function training activities to improve patient motivation and increase engagement with the training interventions. Video gaming has been shown to increase brain dopamine levels^8^, and functional deficits in children with ADHD may improve during video game play^9^. Secondly, the training strategies have expanded to target the training of multiple executive functions, as well as real-life skills requiring the use of executive functions for their application. For example, video game intervention designed to train working memory, inhibition, and cognitive flexibility improved both parent and teacher ratings of ADHD symptoms^10^. Another video game intervention was designed to train real-life skills such as time management, organization, planning, and cooperation, resulting in improvements in both parent and teacher ratings of childrens’ performance in time management (ADHD symptom ratings were not assessed)^11^.

An ongoing clinical pilot study is investigating the use of Minecraft as a therapeutic video game treatment for ADHD in combination with stimulants^12^. Minecraft provides an open-ended video game environment with a considerable degree of flexibility for the development of a variety of executive function training activities. Training activities to address both multiple core executive function deficits and real-life skills resulting from the use of executive functions are included in the present study. In order to address the heterogeneous nature of executive function deficits in ADHD^13^, a personalized treatment intervention that can provide an optimal schedule of executive function training activities for the pattern of deficits present in each individual patient to further improve outcomes is under development. This personalized treatment intervention will also include a stimulant dosage target to reduce the trial and error required to find an optimal dose for a given child with ADHD.

A computational model utilizing artificial neural networks (ANNs) will be used to produce the personalized treatment recommendations. Artificial neural networks are a group of machine learning methods that can learn complex patterns in a dataset and use these patterns to generate predictions for new examples^14^. ANNs have generated a great deal of interest for their applications in healthcare. For example, ANNs have been shown capable of medical imaging diagnostic performance comparable to human physicians^15^, and even outperformed neurologists in diagnosing intracranial hemorrhages from CT scans^16^. Neural network-based technologies have also been developed for use in institutional medical settings to identify early-stage sepsis^17^ and predict the risk of deterioration in intensive care patients^18,19^.

The structure and function of artificial neural networks were inspired by biological neural networks, although they remain a highly simplified approximation. Nonetheless, they can learn to make predictions in complex, changing environments and could therefore be used as a simplified model for the behavior of humans in an environment such as Minecraft. Thus, by developing a set of computational tasks performed by ANNs that resemble executive function tasks performed by human subjects in Minecraft it may be possible to gain insight into how human task performance is optimized by careful inspection of how ANN performance on their computational tasks is optimized. This hypothesis will guide the personalization of the executive function training activity schedule recommendations in the therapeutic video game intervention^20^.

The objective of this work is to develop a computational task performed by ANNs that represents the use of time management skills in humans. This task was designed to require the use of the fluid intelligence and working memory executive functions, with the working memory function impeded by prepotent impulsivity. First, a fluid intelligence function represented by the performance of a factorization of the number 12 by basic recurrent neural networks (RNNs)^21^ was developed. This fluid intelligence function was then combined with the previously developed working memory and prepotent impulsivity functions that used convolutional neural networks (CNNs)^22^ to identify handwritten digits in the MNIST test set^23^ to create the computational Time Management activity.

## Methods and Results

### Generating Fluid Intelligence Deficiencies (Colaboratory Notebook)

Fluid intelligence is the use of adaptive reasoning in novel situations^4^. For the purposes of this project fluid intelligence is considered as a problem solving ability consisting of the mental processes analysis and synthesis^5^. The process of analysis involves the decomposition of complex information into smaller, simpler parts, while the process of synthesis involves the combination of simple pieces of information into more complex, novel combinations. To represent the fluid intelligence process of synthesis in this experiment, a computational Factorization of 12 activity was designed. This activity is performed by RNNs that are trained to produce a sequence of integer outputs in the range 1-10 that can be multiplied together to produce the larger integer 12.

RNNs were chosen to perform the computational Factorization of 12 activity because they are capable of processing sequences consisting of multiple inputs^21^. Although the architectures of RNNs can vary considerably in complexity, the simplest form of RNN possesses a single internal layer that changes its state as the sequence is processed and three sets of weight parameters. One set of weights processes the input to the network, another set of weights modifies the state of the internal layer of the network, and the third set of weights converts the inner state of the network into an output. The inner state of the network changes as each individual element of the input sequence is processed in a series of steps; this inner state is affected by the current element being processed as well as all the previously processed elements and determines the output that the network produces at each step. As a result, the output of the network at each step depends on both the previously processed elements of the sequence and the currently processed element.

To produce a list of integers that are factors of the number 12, the RNN is provided with the start token as an initial input, and outputs a single predicted factor in the first step. This predicted factor is recorded in a list, which is then provided as an input (rather than the start token) for the RNN in the next step (Figure 1). This process repeats until the RNN produces an output indicating the sequence of factors is complete.

**Figure 1 –.**
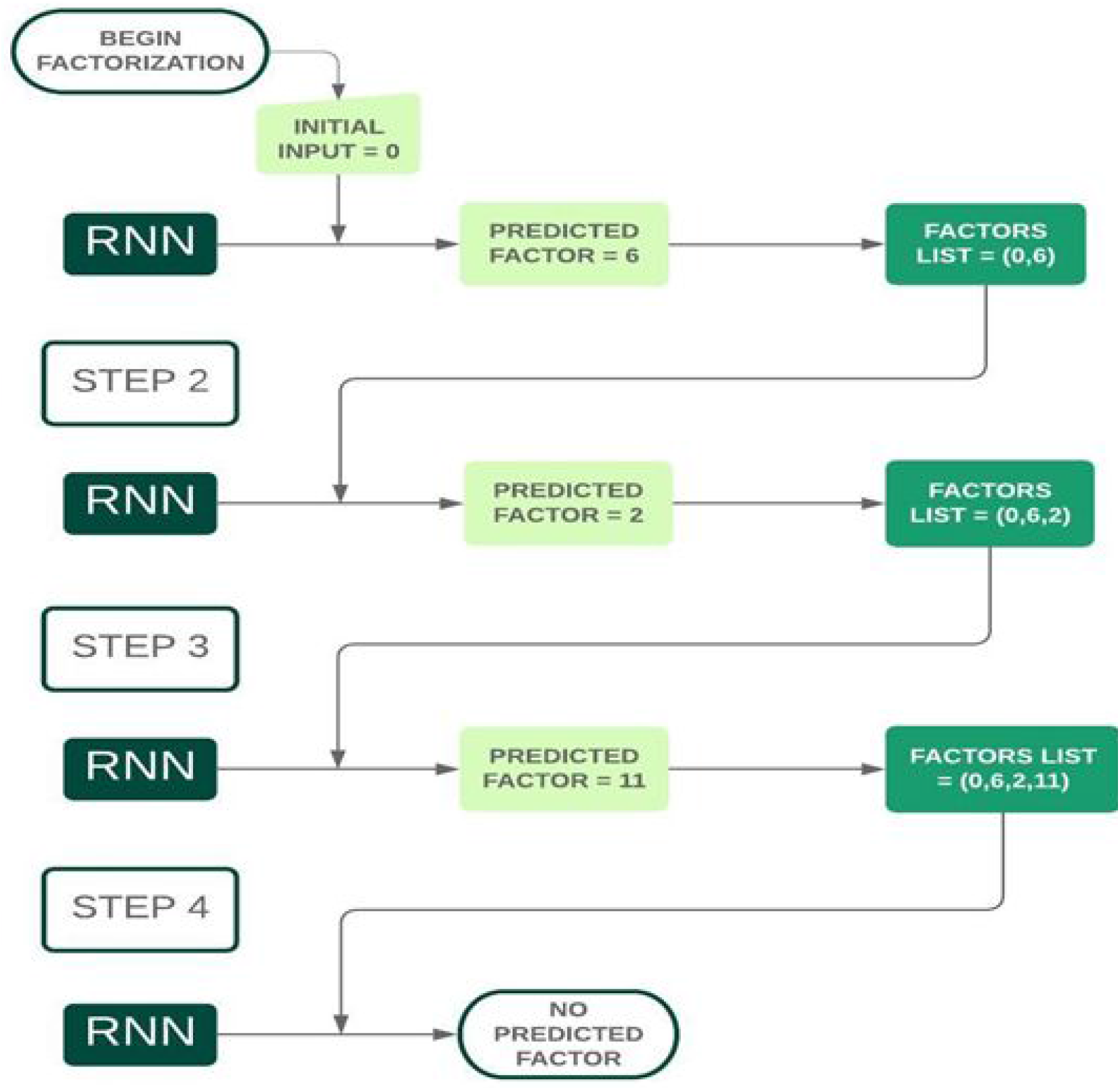
Factor Prediction for the Number 12 by a Recurrent Neural Network

All calculations for this experiment were performed with Python v3 in a Google Colaboratory Notebook. A basic RNN model modified from a RNN designed for language modeling and the generation of reddit comments^24^ was implemented. This RNN contained a single inner layer of 100 units, used the tanh activation function (a non-linear function with outputs ranging from −1 to 1) to calculate the inner layer, and a softmax activation function to generate a list of normalized probabilities for integers in the range 1-11 as an output. The number 0 was used as a “start token”, or initial input to signal the start of a factorization prediction, while the number 11 was used as an “end token” to indicate the end of a sequence of predicted outputs.

A training set composed of 29 training examples with features (inputs) and labels (correct outputs) were generated by hand. The set of training features consisted of lists of integers between 0 and 10, and the list for each training feature began with the integer 0. Lists of integers that could be multiplied together to produce the number 12 and lists of integers that could not be multiplied together to produce the number 12 were both included. The set of training features lists that produced the number 12 included all the permutations of each set of factors, while the training features lists that did not produce the number 12 simply consisted of the start token 0 followed by the incorrect factor. A matching training label was created for each training feature. The training label for each example contained the same list of integers contained in the features for the example, but the start token 0 was removed and the end token 11 was added to the end of the list.

To produce different categories of performance for the computational Factorization of 12 activity, modifications were introduced into a supervised learning training procedure and investigated for their effects on RNN performance. In this case, the data available for training was limited (29 training examples) so the differences in performance were created by varying the number of training cycles (or epochs) as the independent variable. All of the training examples are presented to a neural network in one training epoch, so the number of times a neural network sees the set of training examples is equivalent to the number of training epochs. Ten groups of 12 x RNNs per group were assigned unique durations for their training ranging from 1 training epoch to 5,000 training epochs. The categorical cross-entropy loss function was used to measure the deviation between RNN outputs and correct predictions and calculate gradients to adjust parameter values. Stochastic gradient descent with a learning rate of 0.005 was used for the optimization method to adjust parameter values.

After training, the performance of each RNN was evaluated by calculating its percent accuracy on 100 factorization of 12 predictions. As expected, the mean percent accuracy for the RNNs in each group increased with the number of training epochs in a logarithmic relationship (Figure 2). The minimum group mean accuracy was 0.5 % (SD 0.9%) for the RNN group trained for 1 epoch while the maximum group mean accuracy achieved was 81.1% (SD 2.9%) for the RNN group trained for 5,000 epochs. The rate of improvement in performance began to level off at a group mean accuracy of 74.2% (SD 5.4%) with 250 training epochs, although significant improvements in performance were still observed as the number of training epochs was increased further.

**Figure 2 –.**
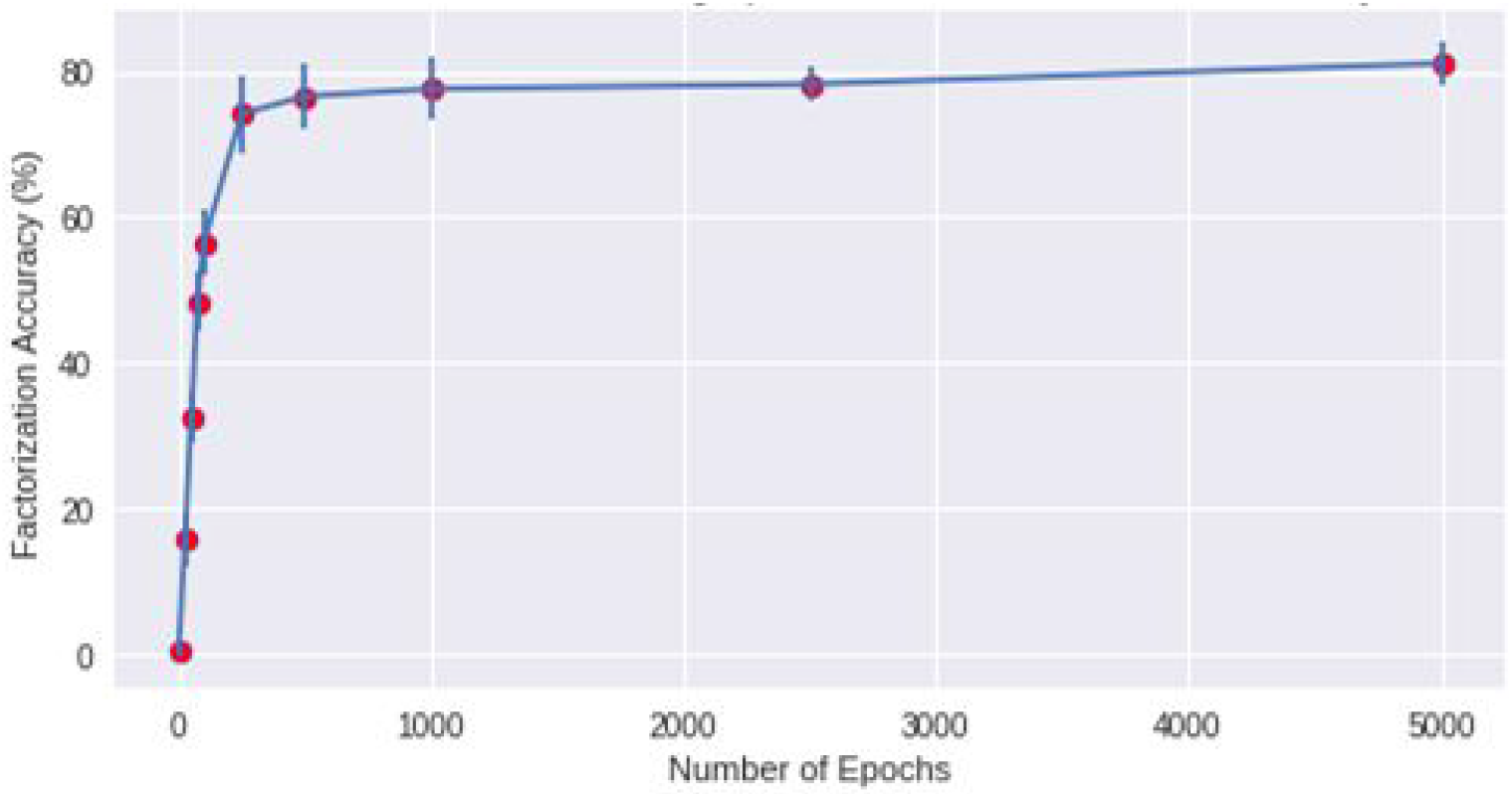
Effect of Number of Training Epochs on Mean Group Accuracy of Factorization of 12 in Recurrent Neural Networks (n = 12 x RNNs Per Group)

The training procedure for “deficient” fluid intelligence RNNs was selected to achieve a goal of approximately 50% mean factorization of 12 accuracy, while the training procedure for “healthy” fluid intelligence RNNs was selected with a consideration of performance and computational efficiency. The goal performance level for “deficient” fluid intelligence was achieved in RNNs trained for 75 epochs (group mean accuracy = 48.3% (SD 4.1%). Training for 1000 epochs was selected as the learning procedure for “healthy” fluid intelligence RNNs to achieve a mean group accuracy of 77.6 % (SD 4.2%) with a sacrifice of 3.5% accuracy in exchange for significantly improved computational efficiency compared to training for 5000 epochs.

In summary, a computational representation for the synthesis function of the higher order executive function fluid intelligence in humans was developed in this experiment. This model includes basic RNNs to predict integers between 1 and 10 that can be multiplied together to produce the number 12, where a “healthy” fluid intelligence is represented by RNNs with high Factorization of 12 accuracy and a “deficient” fluid intelligence is represented by RNNs with poor Factorization of 12 accuracy. This Factorization of 12 fluid intelligence function was then integrated together with the MNIST handwritten digit recognition function representation for working memory and the prepotent impulsivity function to create the first implementation of virtual “subjects” who perform a virtual Time Management task.

### Time Management Task (Colaboratory Notebook)

A pertinent example of the use of fluid intelligence is in the practice of time management. When considering how to complete a complex task most efficiently, one may first analyze the task carefully to determine the individual steps needed for its completion. There will be a finite number of ways to organize these steps, and the best sequence of steps will be the one that produces the best outcome in the shortest amount of time. Working memory and the synthesis process of fluid intelligence can be used to mentally rehearse the completion of varying combinations of these steps and determine the time required to complete them. Some combinations of steps may overlap better in time than others when they are completed sequentially, and some steps may be found to provide minimal contributions to the overall goal. The order of the overlapping steps may then be preserved in the final sequence, and the marginally important steps may be left at the end of the sequence to be completed after the more highly prioritized steps should time permit. In this way, fluid intelligence can work together with the core executive function working memory in a time management activity.

While the working memory and fluid intelligence functions are each represented by the performance of individual ANNs in performing a given activity representing an executive function, the Time Management task consists of a group of ANNs performing their given activities in tandem to complete a given task together. This combination of multiple ANNs working together to complete a task can be thought of as a virtual “subject” that utilizes multiple executive functions to complete a computational representation of an executive function training activity completed by human subjects in the therapeutic video game intervention (Figure 3).

**Figure 3 –.**
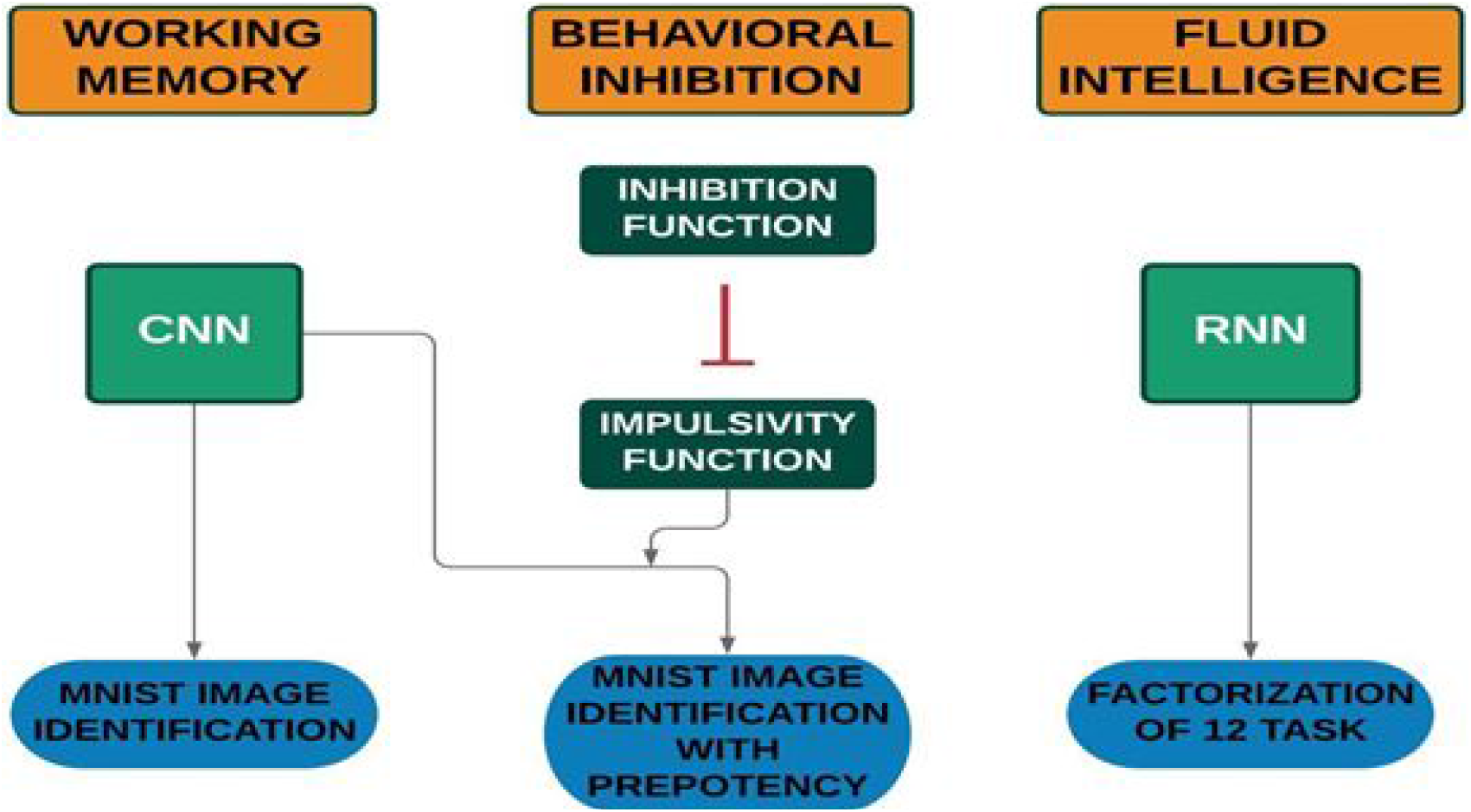
Building the Virtual “Subject”

Similar to the fluid intelligence task described in the previous section, the goal of the Time Management task is to create a list of integers between 1 and 10 that can be multiplied together to produce the number 12. However, the process required to achieve this goal is modified in three important ways. Firstly, the Time Management task is not complete after a single factorization of 12 attempt, but rather continues until a successful factorization of 12 is completed (Figure 4). Secondly, the Time Management task requires the use of the working memory function impeded by the prepotent impulsivity function for its completion. The working memory function with prepotent impulsivity is used after the prediction of a single factor to locate and correctly identify an MNIST image that contains the predicted factor as its handwritten digit in the MNIST test set. Finally, the performance of the Time Management task is measured by the total number of MNIST test set images searched through to complete a successful factorization of 12 rather than the percentage of successful factorizations.

**Figure 4 –.**
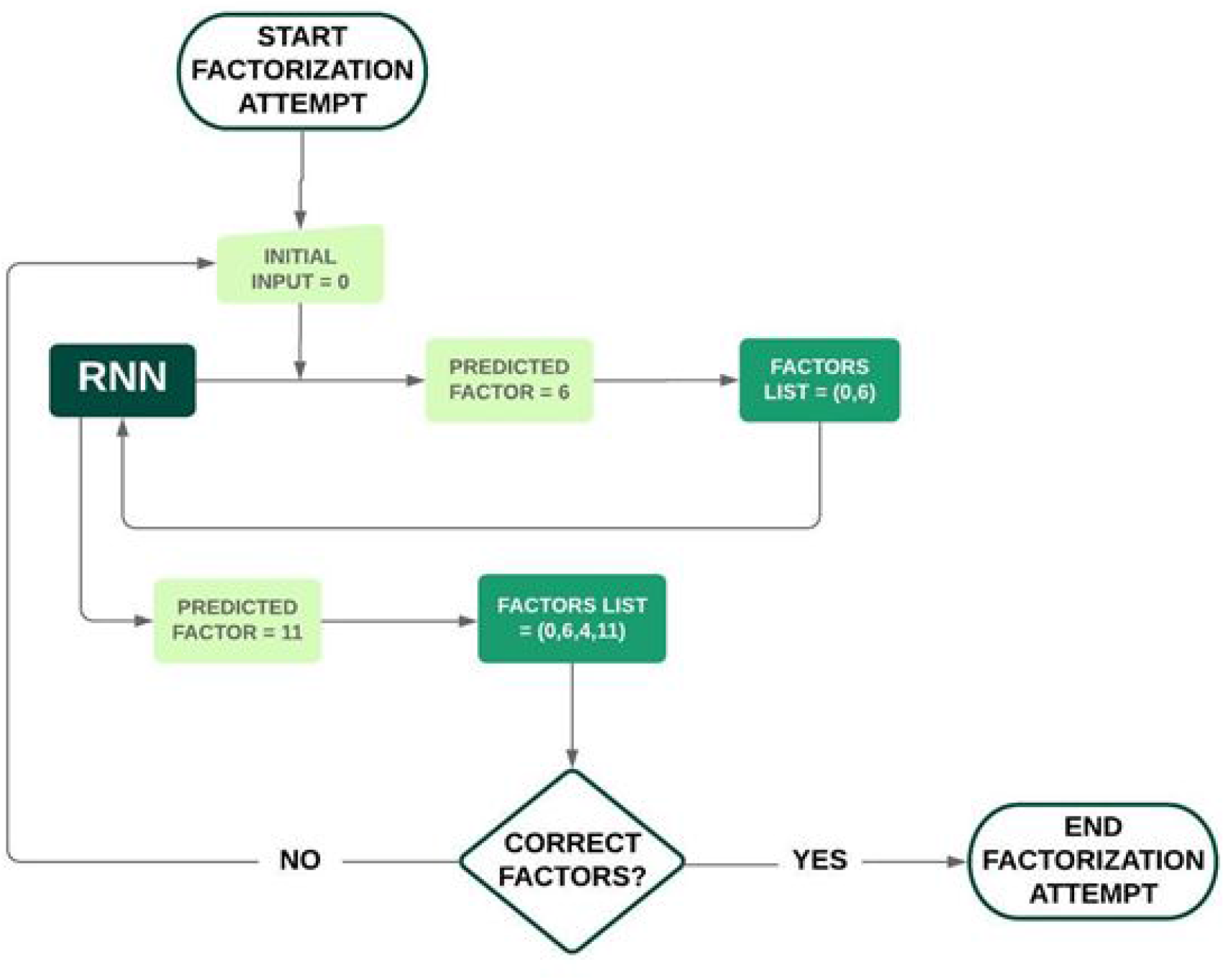
Recurrent Neural Networks (Fluid Intelligence) in the Time Management Task

The procedure to create virtual “subjects” who complete the virtual Time Management task is simply the combining of the previously developed neural network training procedures. Each virtual “subject” consists of a working memory function consisting of a CNN trained for handwritten digit recognition in the MNIST test set, a set of parameters for the prepotent impulsivity function that impedes the performance of the working memory function, and a fluid intelligence function consisting of an RNN trained to factorize the number 12. Two groups of virtual “subjects” with n = 6 x virtual “subjects” per group were created: a control group with “healthy” working memory and fluid intelligence and an “ADHD” group with “deficient” working memory and fluid intelligence. Prepotent impulsivity^25^ was included in all “subjects” in both groups with the control group possessing a set of parameters that leads to lower impulsivity than the “subjects” in the “ADHD” group.

All computations for the Time Management task were performed with Python v3 in a Google Colaboratory Notebook, and the creation and training of CNNs were performed with the Keras Machine Learning Library^26^. Data visualizations were created with R v3.6. ANNs for the virtual “subjects” in the control group were trained with the learning procedures that produced 6x “healthy” working memory and 6x “healthy” fluid intelligence representations, while ANNs for the virtual “subjects” in the ADHD group were trained with the learning procedures that produced 6x “deficient” working memory and 6x “deficient” fluid intelligence representations. All ANNs (CNNs^25^ and RNNs) for each executive function representation possessed an identical architecture across both groups as previously described. “Healthy” working memory representations for six virtual control “subjects” were produced by training each of six CNNs with 2,500 handwritten digit images randomly selected from the MNIST training set and “deficient” working memories for six virtual “ADHD subjects” were produced by training each of six CNNs with 25 randomly selected MNIST training set images. Similarly, “healthy” fluid intelligence representations for six virtual control “subjects” were produced by training each of six basic RNNs for 1,000 epochs to factorize the number 12 and “deficient” fluid intelligence representations for six virtual “ADHD subjects” were produced by training each of six basic RNNs for 75 training epochs. Baseline accuracies of the working memory representations were measured by handwritten digit recognition by CNNs on the MNIST test set while baseline accuracies of the fluid intelligence representations were determined on 100x factorization of 12 attempts by RNNs. The Mann-Whitney U test was used to test for significant differences between group means in baseline performance for each executive function representation.

After training and evaluation of baseline performance of all ANNs, the Time Management task was performed by the virtual “subjects” (Figure 5). In the first step of the task, the virtual “subject’s” fluid intelligence representation uses a basic RNN to perform a single factor prediction. The identity of this factor is then provided to the virtual “subject’s” working memory representation with prepotent impulsivity, and the virtual “subject” begins to identify handwritten digits in randomly selected images from the MNIST test set. The virtual “subject’s” prepotent impulsivity function is operational with each identification, and may activate an automatic impulsive response where the virtual “subject” simply responds with its previous handwritten digit recognition output rather than using its working memory representation to identify the handwritten digit with a CNN. The automatic impulsive response becomes more likely as multiple MNIST images containing the same handwritten digit are encountered in succession. This search continues until the predicted factor is located AND correctly identified in the MNIST test set. Once the search is successfully completed, another factor prediction is performed and this sequence of steps continues until a complete factor list is predicted and all of its elements are correctly identified in the MNIST test set. When a correct factor list is predicted one trial of the task is complete. Otherwise, this procedure is repeated until a correct factor list is produced. Each virtual “subject” completed 25 successful factorizations of 12 trials, and a mean MNIST test set images searched count for all the trials was calculated for each “subject”. The Mann-Whitney U Test was then used to test the group mean images searched counts for significant differences.

**Figure 5 –.**
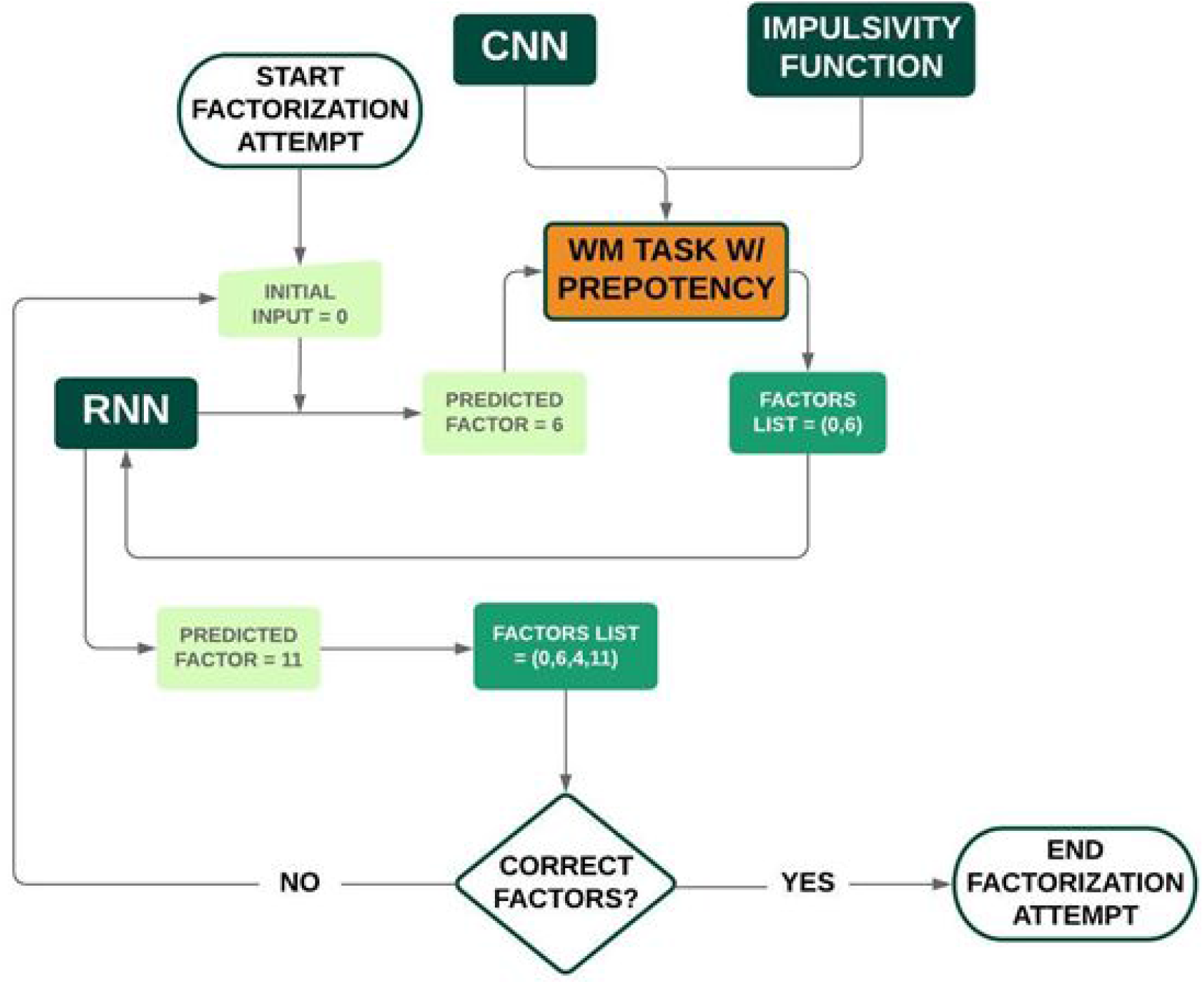
Time Management Task Flowchart

Without prepotent impulsivity, virtual “subjects” in the “healthy” control group achieved a baseline group median handwritten digit recognition accuracy of 95.9% (IQR 0.6%) while virtual “subjects” in the “ADHD” group achieved a baseline group median handwritten digit recognition accuracy of 40.2% (IQR 7.2%) on the MNIST test set (Figure 6). A Mann-Whitney U Test showed a significant difference between the groups (p = 0.025).

**Figure 6 –.**
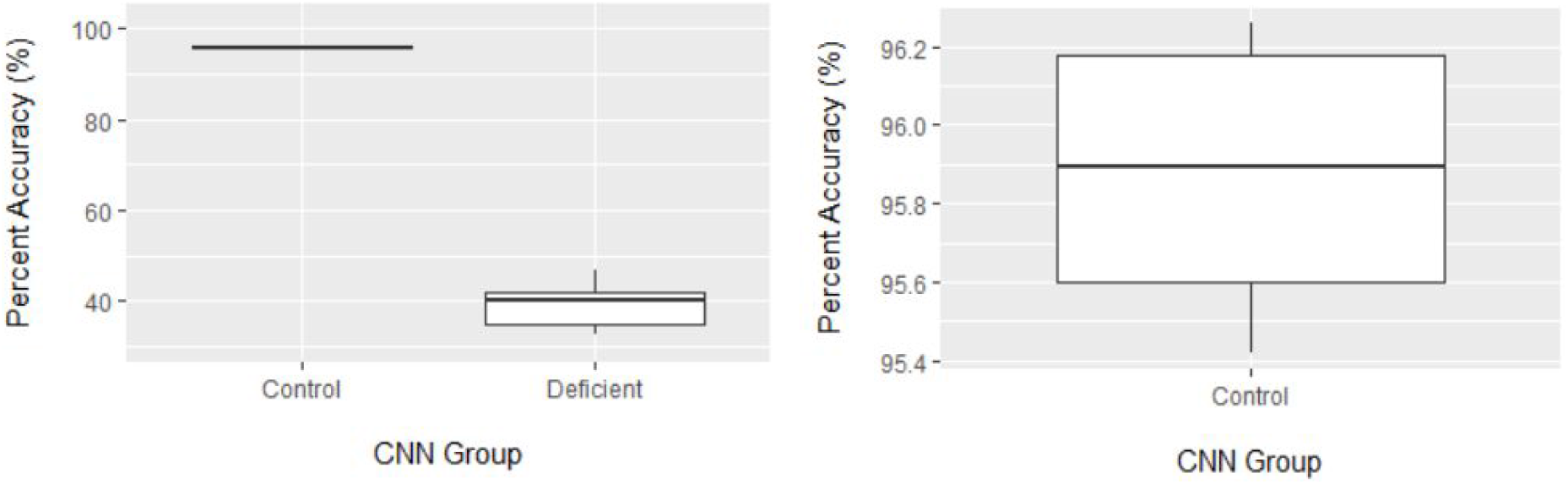
Baseline Difference in Mean Group Accuracy of Handwritten Digit Recognition on MNIST TestSet by Convolutional Neural Networks of Virtual “Subjects” in the Time Management Task (n = 6 x CNNs Per Group)

Virtual “subjects” in the “healthy” control group achieved a baseline group median factorization of 12 accuracy of 78.5% (IQR 6.0%) while virtual “subjects” in the “ADHD” group achieved a baseline group median factorization of 12 accuracy of 46.5% (IQR 5.5%) (Figure 7). A Mann-Whitney U Test revealed a significant difference between groups (p = 0.025). Taken together, these data indicate that a significant difference in performance of both the working memory and fluid intelligence functions were successfully produced between the virtual “subjects” in the “healthy” control and “ADHD” groups.

**Figure 7 –.**
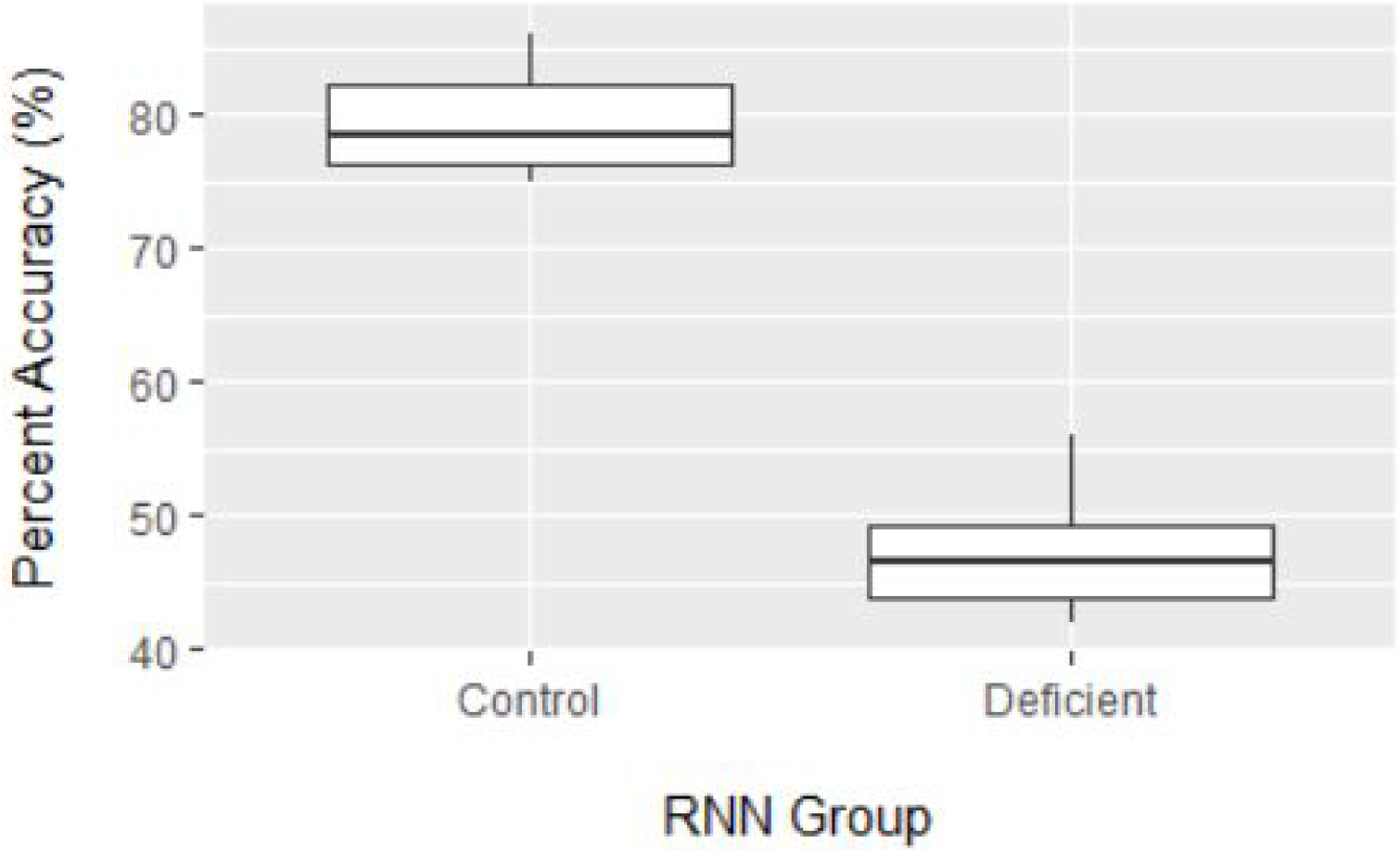
Baseline Difference in Mean Group Accuracy of Factorization of 12 by Recurrent Neural Networks of Virtual “Subjects” in the Time Management Task (n = 6 x RNNs Per Group)

Finally, virtual “subjects” in the “healthy” control group required a group median of 31.5 (IQR 13.3) MNIST test set images searched through, while virtual “subjects” in the “ADHD” group required a group median of 282.5 (IQR 331.3) MNIST test set images searched through to complete the virtual Time Management task (Figure 8). A Mann-Whitney U Test showed a significant difference between groups (p = 0.0025). These results indicated that this model composed of virtual “subjects” built from executive function representations of working memory and fluid intelligence with impulsivity that complete a virtual Time Management task that requires the use of these executive function representations was able to produce two groups of “subjects” with distinct levels of performance in the virtual task.

**Figure 8 –.**
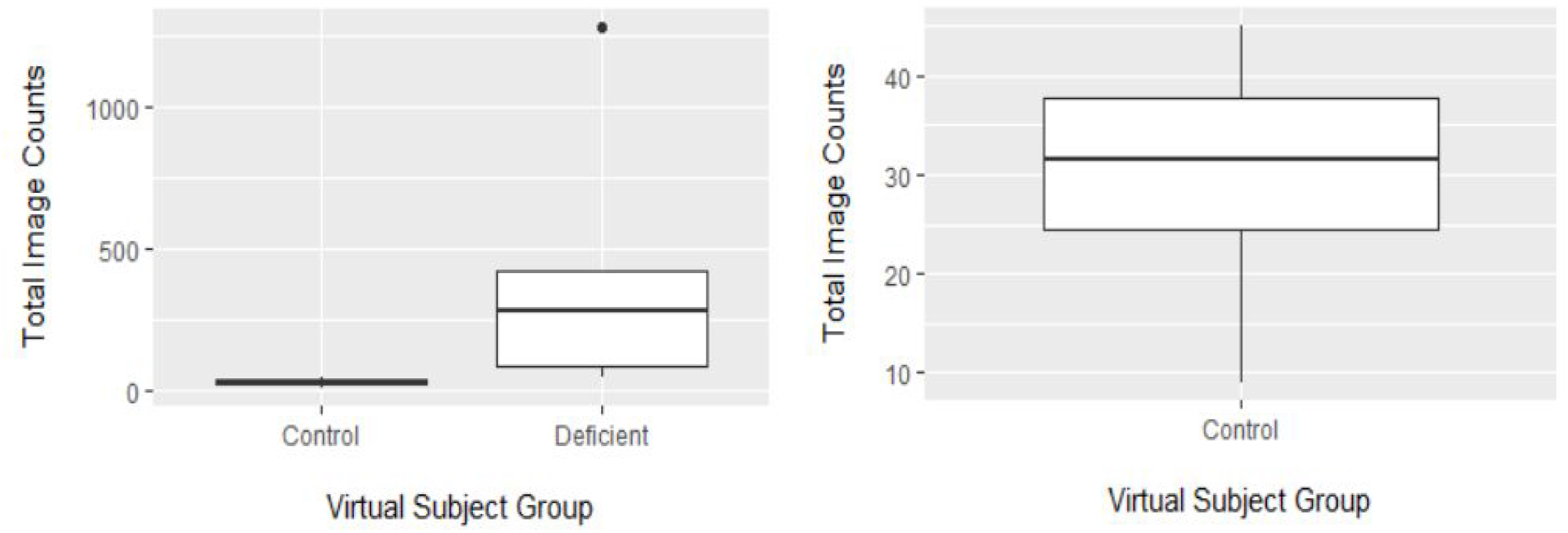
Performance of Virtual “Subjects” in the Time Management Task (n = 6 x Virtual “Subjects” Per Group)

## Discussion

In summary, this experiment used a basic RNN model to serve as a simplified model of fluid intelligence, and a procedure was developed to produce “healthy” and “deficient” fluid intelligence performance. This fluid intelligence representation was combined with the previously developed working memory & impulsivity representations to create a virtual “subject” who completed a simulated Time Management task that required the use of these executive function representations. Finally, a significant performance difference between “healthy” and “deficient” virtual “subjects” in this task was demonstrated.

The Factorization of 12 task using basic RNNs to output factor sets of the number 12 was used as a representation for the synthesis component of fluid intelligence in this experiment. This task can be improved by the inclusion of a representation for the process of analysis to create a more comprehensive model of fluid intelligence and by adopting a different variety of RNN to increase the complexity of the task. Consider the process of analysis (breaking down complex ideas) as the inverse of the process of synthesis (building up complex ideas). When the process of synthesis is then represented as the building up of the “complex” number 12 from its simpler component factors, the process of analysis may be represented as the breaking down of the number 12 into its simpler component factors. Thus, the fluid intelligence operation that currently consists of the process of synthesis will be expanded to include an RNN task to break down the number 12 as a representation for the process of analysis.

In the current version of the task, the RNN begins by producing a factor of the number 12, which is then used as an input in the next loop of the task for the RNN to output another factor. This process continues until the RNN produces an output to indicate the end of the sequence of factors and then concludes with an evaluation of the accuracy of the sequence. The reverse process can be implemented through the modification of the RNN’s training set. The training set will be differentiated into two separate categories, where one category consists of factor sequences with factors ordered from smallest to largest in size and vice-versa for the other category. The training set with examples ordered by increasing size of their factors will train RNNs for the process of synthesis (where the larger number is assembled), while the training set with examples ordered by decreasing size will train RNNs for the process of analysis (where the larger number is broken down).

The fluid intelligence model can be improved by the substitution of the basic RNNs with Long-Short Term Memory networks (LSTMs)^27^. While basic RNNs indiscriminately update every parameter of their internal layer with each step of the sequence they produce, LSTMs include additional “forget” gates in this updating process that screen for and exclude the passage of extraneous information to the internal layer between steps in the sequence. This increased efficiency of updating permits the processing of larger sequences in LSTMs, and thus LSTMs would be beneficial to permit the analysis and synthesis of larger numbers in this project. The capability to process larger numbers in this way would allow the creation of multiple difficulty levels for the task to more accurately capture the properties of the therapeutic video games tasks which have multiple difficulty levels.

A comparison of this representation of fluid intelligence to alternative modeling approaches is difficult due to a paucity of research related to the neural functions that produce this executive function. The function of a network of brain regions known collectively as the Multiple-Demand System has been proposed to play a general role across multiple complex problem solving tasks in primates^28^. Detailed functional neural network theories are lacking however. While a recent theory has proposed that fluid intelligence results from a widespread, randomly connected network structure of groups of neurons^29^, to our knowledge there does not seem to be any research available that examines the computations that these types of networks conduct. However, it is recognized that this representation of fluid intelligence is grossly simplified compared to the underlying biological functions. Nonetheless, an accurate representation of the functional complexity of fluid intelligence may not be a necessary prerequisite for modeling the effects of the learning processes that improve its performance.

In addition to their potential applications in drug discovery and the practice of medicine, ANNs and other machine learning methods have also drawn interest as tools to improve clinical trial execution and design. For example, the patient selection process can be aided by search tools that are able to interpret and synthesize patient health information from a variety of sources, and patient adherence & retention can be improved by wearable technologies that report data from clinical monitoring parameters to machine learning systems that then analyze it^30^. Furthermore, machine learning models are under development to predict disease progression, such as in Alzheimer’s disease^31^. These models can be used to inform both patient selection and clinical trial design to increase the likelihood that a given clinical endpoint can be achieved successfully^32^. This work also aims to enhance clinical trial design, but does so in a unique way. Rather than modelling the disease process to optimize the effects of a single treatment during a clinical study, this simulation seeks to model the effects of a variety of treatment options to inform the selection of an optimal intervention.

The Time Management task described here is the first of a series of computational representations of executive function training activities that are part of a therapeutic video game treatment utilizing the online platform Minecraft to treat children with ADHD. This intervention will utilize a novel personalized medicine approach where an individualized treatment regimen consisting of an initial stimulant dose recommendation and schedule of therapeutic video game activities will be determined from the initial ADHD assessment results of new patients. We hypothesize that the training of ANNs can serve as a computational model for the adaptations that biological neural networks undergo in response to practice of activities requiring the use of executive functions. In the model, a set of ANNs will first be trained to resemble the pattern of executive function deficits reflected in a new patient’s assessment responses. A large set of virtual tasks resembling the therapeutic video game activities will be tested in the model, and the set of virtual tasks that produces the maximum improvements in the ANNs of the virtual “subject” will predict the analogous set of therapeutic video game tasks most beneficial for the human patient. A future clinical study will test this hypothesis by including an experimental group with a personalized executive function activity training schedule guided by the computational model that will be compared with a control group completing an executive function activity training intervention that is not guided by the computational model.

